# Losartan controls immune checkpoint blocker-induced edema and improves survival in glioblastoma

**DOI:** 10.1101/2022.06.28.497997

**Authors:** Meenal Datta, Sampurna Chatterjee, Elizabeth M. Perez, Simon Gritsch, Sylvie Roberge, Mark Duquette, Ivy X. Chen, Kamila Naxerova, Ashwin S. Kumar, Mitrajit Ghosh, Kyrre E. Emblem, Mei R. Ng, William W. Ho, Pragya Kumar, Shanmugarajan Krishnan, Xinyue Dong, Maria C. Speranza, Martha R. Neagu, David A. Reardon, Arlene H. Sharpe, Gordon J. Freeman, Mario L. Suvà, Lei Xu, Rakesh K. Jain

## Abstract

Immune checkpoint blockers (ICBs) have failed in all Phase III glioblastoma trials. Here, we found that ICBs induce cerebral edema in some patients and mice with glioblastoma. Through single-cell RNA sequencing, intravital imaging, and T cell blocking studies in mice, we demonstrated that this edema results from an inflammatory response following anti-PD1 antibody treatment that disrupts the blood-tumor-barrier. Used in lieu of immunosuppressive corticosteroids, the angiotensin receptor blocker losartan prevented this ICB-induced edema and reprogrammed the tumor microenvironment, curing 20% of mice which increased to 40% in combination with standard of care treatment. Using a bihemispheric tumor model, we identified a “hot” tumor immune signature prior to losartan+anti-PD1 therapy that predicted long-term survival. Our findings provide the rationale and associated biomarkers to test losartan with ICBs in glioblastoma patients.

**One-Sentence Summary:** Losartan prevents immunotherapy-associated edema and enhances the outcome of immunotherapy in glioblastoma.

## Main Text

Despite reports that some murine glioblastoma (GBM) models can be cured with immune checkpoint blockers (ICBs), this immunotherapeutic approach has failed in all phase III GBM clinical trials. A challenge unique to GBM is the cerebral edema which can be exacerbated by anti-programmed death/ligand 1 (PD1/PD-L1) antibodies (*1, 2*). Currently, this increased edema can be controlled by potent, immunosuppressive steroids that compromise ICB efficacy.

Here, we demonstrate that the angiotensin receptor blocker (ARB) losartan prevents ICB-induced edema by reducing tumor endothelial cell (TEC) expression of membrane-type matrix metalloproteinases 1 and 2 (MT-MMP-1, -2) that are upregulated during treatment with an anti-PD1 antibody. Furthermore, losartan increases GBM perfusion, enhances anti-tumor immunity, and improves survival (in 2 out of 3 models) under anti-PD1 treatment. Utilizing a bihemispheric model, we show that TME immune composition prior to treatment predicts individual and differential response.

## Results

### ICB treatment disrupts the GBM vasculature and induces edema

MR imaging revealed ICB-induced edema in some GBM patients (**Fig 1A,B**). In the GL261 model, anti-PD1 antibody treatment recapitulated this increased edema (**Fig. 1C**). We performed intravital microscopy after injecting the mice with a fluorescent tracer to detect vascular leakage. We found that tumor vessels in control (IgG-treated) mice retained most of the tracer (**Fig. 1D**), but in anti-PD1-treated mice (**Fig. 1E**), excess tracer leaked into the surrounding tissue (**Fig. 1F**), indicating endothelial barrier disruption. Because losartan and other ARBs have been shown to lower vascular endothelial growth factor (VEGF) expression in GBM models and vasogenic edema in retrospective patient studies (*3-5*), we decided to test the effects of losartan treatment on ICB-induced edema.

**Figure 1.**
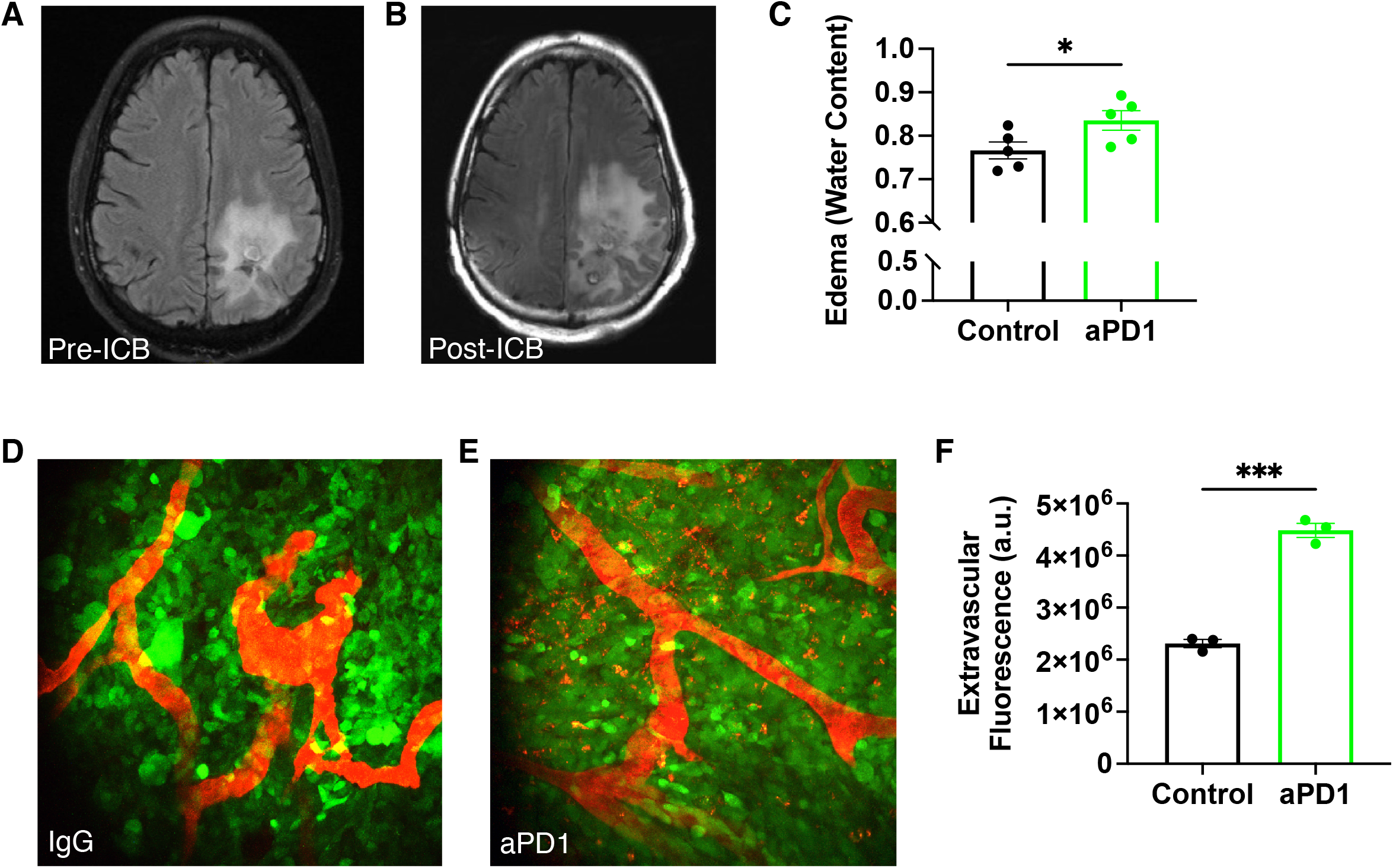
ICB increases GBM vascular leakage and induces brain edema. MR T2-FLAIR images obtained from a recurrent GBM patient (**A**) before and (**B**) after 4 months of anti-PD-L1 (MEDI4763; NCT02336165) treatment show increased edema after ICB treatment. In addition to ICB-induced inflammation, this change may be due in part to underlying tumor activity or growth. (**C**) In mice, anti-PD1 antibody (aPD1) treatment increases edema in GL261 tumors compared to IgG control (as measured by wet-dry weight (i.e., water content) evaluation of tumor tissue; n=5). Multiphoton visualization of the brain vasculature via injected tetramethylrhodamine (TAMRA) labeled albumin (red) imaged through transparent cranial windows in mice bearing GFP+ GL261 GBM (green) shows that compared to IgG controls (**D**) there is increased extravasation in anti-PD1-treated tumors after the third consecutive dose (**E**). (**F**) Quantification shows that more albumin in anti-PD1-treated mice has leaked outside of the tumor blood vessels (n=3). (*Bar plots: mean±SEM; Student’s unpaired t-test; * = p<0*.*05; *** = p<0*.*001*.)

### Losartan prevents ICB-induced edema by reducing TEC MT-MMP-1 and -2 expression

In the GL261 and 005 GSC models (**Fig. 2A, B**), but not in CT2A (**Fig. 2C**), we found that anti-PD1 treatment increased edema, while losartan prevented this anti-PD1-induced edema. To reveal the edema-reduction mechanism, we performed single cell RNA sequencing (scRNASeq) on TECs in the GL261 model (**Fig. S1; Data S1**). We identified a set of genes downregulated in TECs from losartan+anti-PD1-treated tumors vs. anti-PD1 monotherapy (**Fig. 2D,E, Data S2**). This edema signature was most highly expressed in TECs from anti-PD1-treated tumors (**Fig. 2F,G**). Genes included those related to metabolism, angiogenesis/migration, solute carriers, and most notably, a specific subset of MT-MMPs (*Mt1* and *Mt2*, i.e., MMP14 and MMP15). Interestingly, we did not observe gene expression changes in VEGF/VEGFRs or other known vasogenic edema-related genes in this TEC signature (**Fig. 2D,E**). Thus, we explored possible inflammatory mechanisms governing ICB-induced edema.

**Figure 2.**
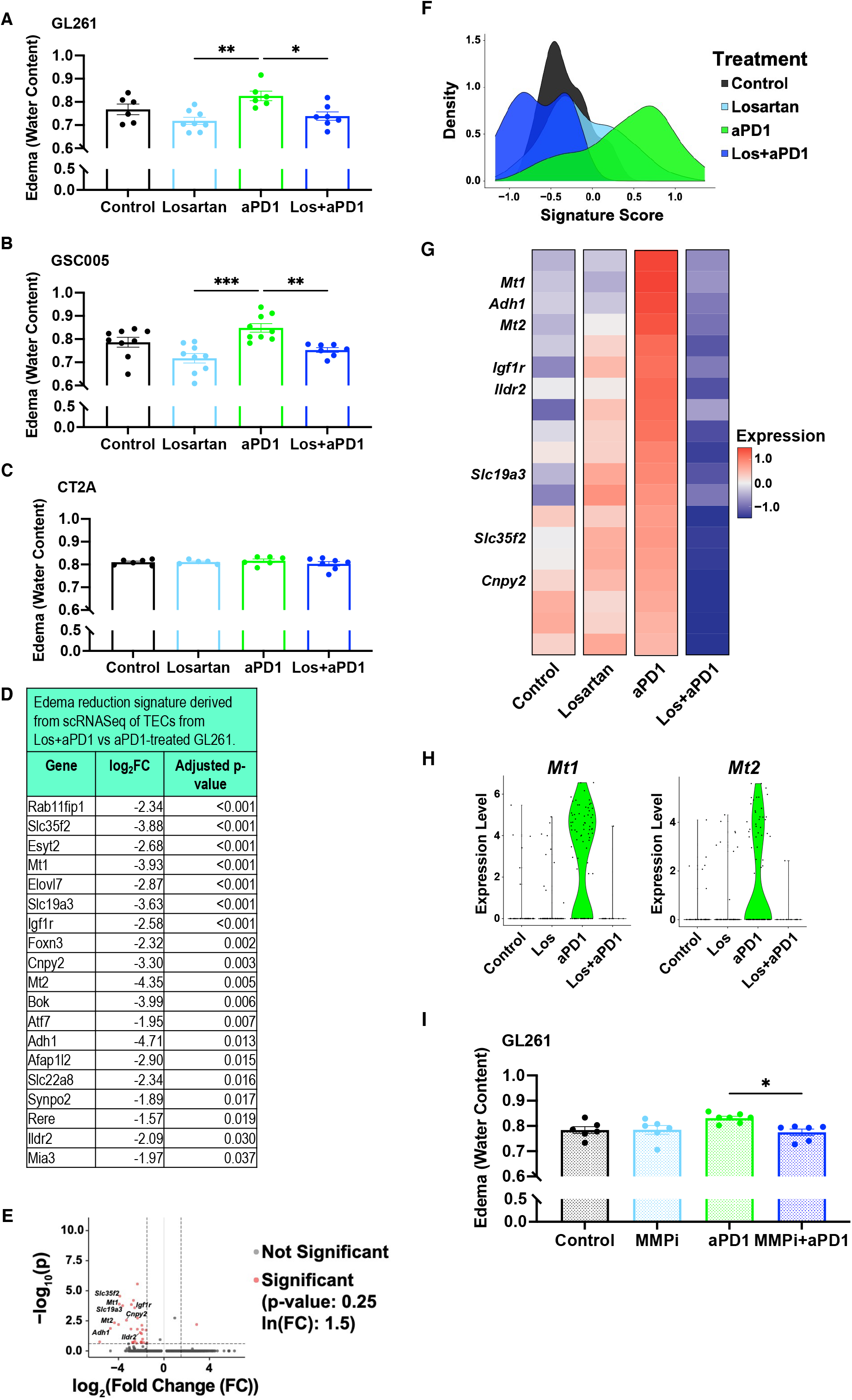
Losartan prevents ICB-induced edema by downregulating TEC MT-MMP-1 and -2 expression. Losartan decreases anti-PD1-induced edema in (**A**) GL261 and (**B**) 005 GSC models but not in (**C**) CT2A after 2 weeks of treatment (n=5-9). (**D**) scRNASeq of TECs reveals a set of downregulated genes that includes those related to metabolism (e.g., *Adh1, Ildr2*), angiogenesis/migration (e.g., *Cnpy2, Igf1r*), solute carriers (e.g., *Slc35f2, Slc19a3*). When applied as an edema signature, this gene set is upregulated in anti-PD1-treated GL261 tumors compared to other treatment arms as visualized via (**E**) volcano plot, (**F**) density plot of edema signature scores (methods described in Supplementary Materials) by treatment and (**G**) mean gene expression heat map of edema signature genes. (**H**) Specialized MT-MMPs (*Mt1, Mt2*) are among these genes and are expressed in TECs only from the anti-PD1-treated tumors. (**I**) The MMP-inhibitor Ilomastat (MMPi) controls anti-PD1-induced edema comparably to losartan in GL261 (n=6). (*Edema signature*: *gene expression units = ln(TP100k +1); log*_*2*_*FC = fold changes>*|*2*|; *adjusted p-value < 0*.*05. Bar plots: mean±SEM; one-way ANOVA with Tukey’s post-hoc test; * = p<0*.*05; ** = p<0*.*01; *** = p<0*.*001*.)

We found via scRNASeq (**Fig. S2; Data S3**) and T cell blocking experiments (**Fig. S3**) that CD8 T cells are important mediators of ICB-induced edema. Because MMP overexpression in endothelial cells has been linked to blood-brain-barrier (BBB) tight junction disruption and cerebral edema (*6, 7*), and can be induced by T cell interactions (*8*), we hypothesized that this could be a potential mechanism of ICB-induced edema in GBM. Indeed, *Mt1* and *Mt2* are only expressed in TECs from anti-PD1-treated tumors (**Fig. 2H**). To test this mechanism, we gave Ilomastat, a broad spectrum MMP-inhibitor that is non-toxic to GBM cells at physiological levels (*9*), to mice bearing GL261 tumors under anti-PD1 treatment. We found that Ilomastat phenocopied the ability of losartan to prevent anti-PD1-induced edema (**Fig. 2I**). Because ARBs can modulate other TME features (*10-13*), we next evaluated the effects of losartan on GBM extracellular matrix (ECM), vasculature, and immune components.

### Losartan reduces ECM and solid stress, normalizes the tumor vasculature, improves perfusion, and decreases hypoxia and immunosuppression in GBM

Losartan lowers collagen and hyaluronic acid (HA) levels in extracranial tumors, reducing the physical force “solid stress,” thereby decompressing previously collapsed blood vessels (*11*). Using bulk RNASeq in GL261, we found that losartan treatment significantly reduced gene expression related to ECM, angiogenesis, immunosuppression and hypoxia compared to controls (**Fig. 3A,B**). We observed reduced expression of immune checkpoints both at the transcriptional (**Fig. 3B**) and protein (**Fig. S4**) levels. Because HA is a major GBM ECM component, we confirmed via immunohistochemistry that losartan lowers HA levels (**Fig. S4**). To test if this reduced solid stress, we analyzed tumor tissue deformation (i.e., a measure of solid stress (*14*)), and found a reduction in losartan-treated tumors (**Fig. S4**).

**Figure 3.**
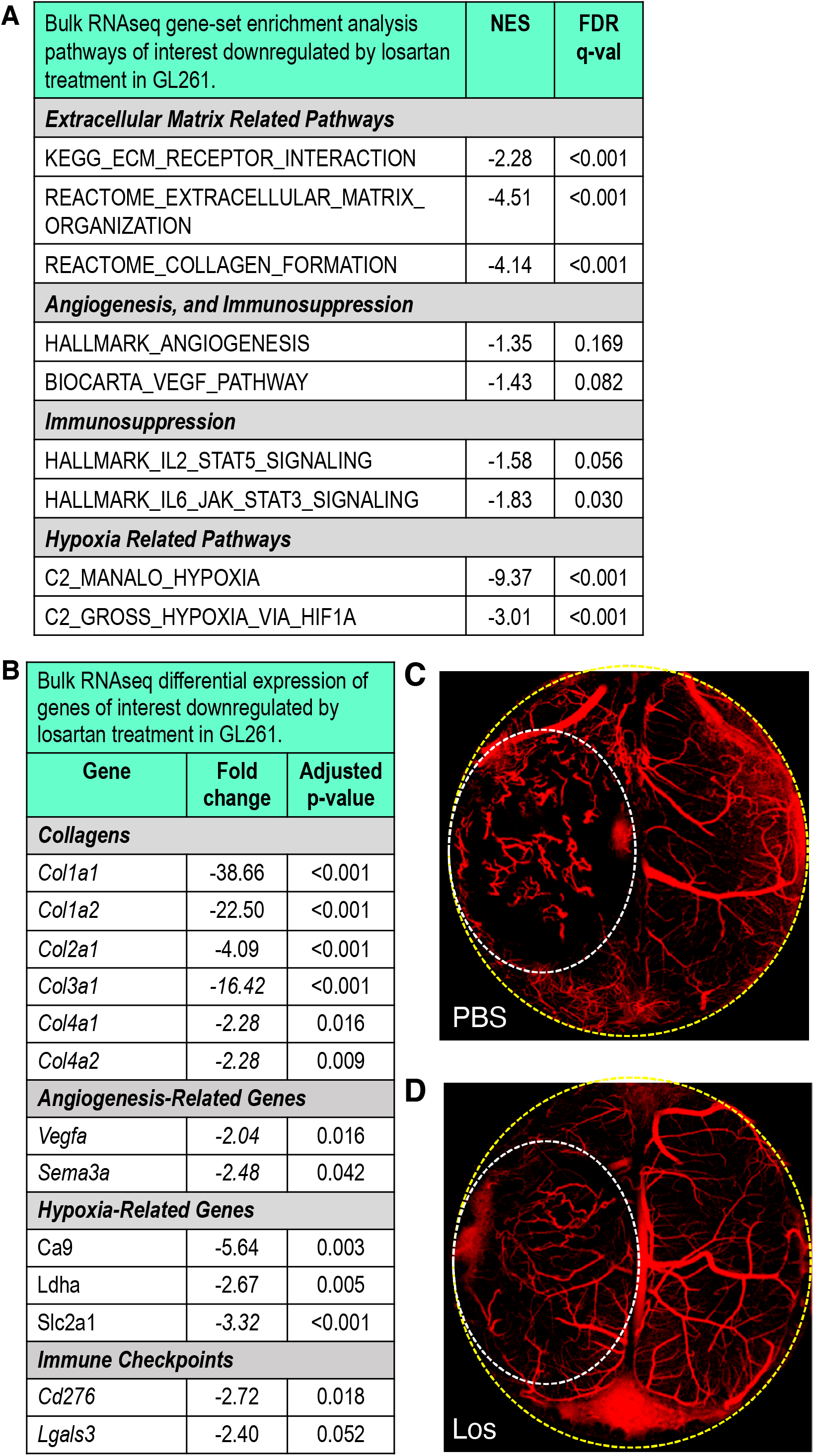
Losartan reprograms the glioblastoma tumor microenvironment. (**A**) TME-related gene-set enrichment analysis pathways downregulated by losartan treatment compared to control in bulk RNASeq of GL261 tumors (n=3). (**B**) Differential gene expression confirms these effects in matrix molecules such as collagen, hypoxia-related genes, and immune checkpoints. Intravital OCT imaging (to detect perfused vessels [red] vs. non-perfused areas [black]) shows that compared to PBS-treated controls (**C**), losartan (**D**) renders tumor blood vessels less tortuous and improves tumor perfusion (yellow dashed line – cranial window border; white dashed line – tumor area). (*Sequencing*: FDR – *false discovery rate*; *all FDR q-values<0*.*20; NES – Normalized Enrichment Score; all adjusted p-values<0*.*05, FC>*|*2*|. *Bar plots: mean±SEM; Student’s unpaired t-test; * = p<0*.*05*.)

We next determined if losartan improved vascular function in GBM. Using optical coherence tomography (OCT) (*15*), we found that control tumors featured chaotic abnormal vessels and non-perfused regions (**Fig. 3C; Movie S1**), whereas losartan-treated tumors had more normalized, straighter, decompressed vessels with greater overall perfusion (**Fig. 3D; Movie S2**). In perfusion-MR images, we found that GBM patients receiving losartan or other angiotensin system inhibitors also had improved tumor perfusion (**Fig. S5**).

### Losartan repolarizes myeloid cells from pro-to anti-tumor phenotype in GBM

To further explore the beneficial mechanisms of losartan on the TME, we next examined tumor-associated macrophages (TAMs) and resident microglia, as both human and murine GBMs are highly infiltrated by these cells. From bulk RNASeq analyses, we found that losartan upregulated microglia-associated genes (**Fig. 4A**), and reduced the expression of global (**Fig. 4A**) and pro-tumor (“M2-like”) TAM-associated genes (**Fig. 4B**).

**Figure 4.**
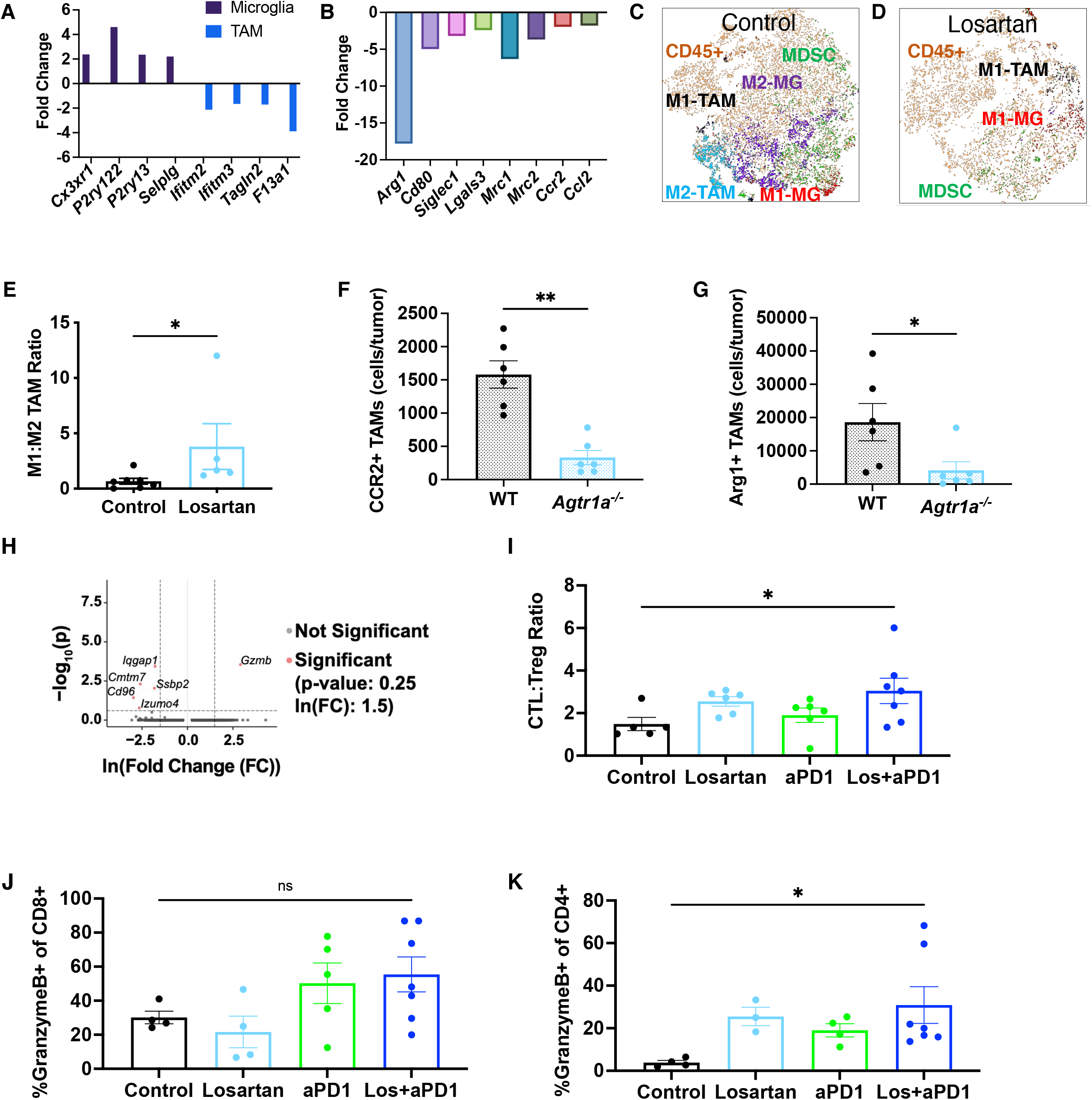
Losartan promotes anti-tumor immunity in the GBM TME. Applying the human-derived signatures from our previous work (*47*), losartan is found to enrich microglia-like signatures and downregulate global (**A**) and “M2-like” (**B**) TAM signatures vs. controls as assessed in bulk RNASeq samples from GL261 (n=3). t-SNE plots of flow cytometry data of myeloid populations reveal (**C**) a diverse and largely immunosuppressive (“M2”) microenvironment in GL261 controls that is (**D**) reprogrammed by losartan treatment to feature fewer myeloid cells that are polarized for anti-tumor (“M1”) activity (MG – microglia). (**E**) Losartan increases the ratio of anti-to pro-tumor TAMs, assessed via flow cytometry (n=5-7). Highly suppressive TAM subsets (**F**) CCR2+ and (**G**) Arg1+ (of CD45hiCD11b+F4/80+) are downregulated in GL261 tumors implanted in *Agtr1a*^*–/–*^ mice compared to those implanted in wildtype (WT) C57Bl/6 mice. (**H**) scRNASeq of CD8+ T cells reveals heightened *Gzmb* expression under combined treatment compared to anti-PD1 monotherapy. Losartan+anti-PD1 treatment increases (**I**) cytotoxic (CTL; CD45+CD3+CD8+GranzymeB+) to regulatory (Treg; CD45+CD3+CD4+FoxP3+) T cell ratios in the tumor, and effector Granzyme+ CD8 (**J**, not significant) and CD4 (**K**) T cells in the cervical lymph nodes. (*Sequencing: all FDR q-val<0*.*25, FC>*|*2*|, *adjusted p-values<0*.*05. Flow cytometry: Mann-Whitney unpaired t-test or one-way ANOVA with Tukey’s post-hoc test; * = p<0*.*05*.)

Using flow cytometry, we found fewer myeloid cells in losartan-treated tumors with reduced M2-like TAM, microglia, and myeloid-derived suppressor cell (MDSC) compartments (**Fig. 4C,D**), and an increased ratio of anti-/pro-tumor (“M1-like/M2-like”) TAMs (**Fig. 4E**). Moreover, pro-tumor TAM populations were significantly reduced in angiotensin type 1 receptor knockout (*Agtr1a*^*–/–*^, i.e., the molecular target of losartan) mice (**Fig. 4F,G**).

### Losartan enhances effector T cell function in GBM during ICB therapy

Based on the ability of losartan to repolarize the myeloid compartment, we next tested the effects of losartan on T cell function in the face of ICB. We found via scRNASeq that CD8 T cells from losartan+anti-PD1-treated tumors had higher expression of *Gzmb* compared to anti-PD1 monotherapy (**Fig. 4H**). By flow cytometry, we found a significantly increased ratio of cytotoxic Granzyme B+ CD8 T cells to regulatory FoxP3+ CD4 T cells during combined losartan+anti-PD1 treatment (**Fig. 4I**), as well as an increase in the overall percentages of granzyme B+ effector T cells (CD8, **Fig. 4J**, and CD4, **Fig. 4K**) in the draining cervical lymph nodes.

Collectively, our results suggest that losartan can reprogram the GBM TME from immunosuppressive to immunostimulatory. Thus, we next explored the ability of losartan to enhance survival under ICB therapy.

### Losartan enhances ICB efficacy without or with the standard of care

Based on the beneficial TME effects of losartan, we designed our survival studies to administer losartan 7 days prior to and throughout anti-PD1 treatment (**Fig. 5A**). In GL261 and 005 GSC models, we found that losartan+anti-PD1 antibody doubled animal survival over anti-PD1 monotherapy, and ∼20% of the mice survived long-term and rejected subsequent tumor re-challenge (**Fig. 5B-E**). However, in the CT2A model (**Fig. 5D**), we observed only a modest benefit of anti-PD1 therapy; adding losartan failed to further enhance ICB efficacy. This is not unexpected, given that CT2A has higher ECM content (**Fig. S4**), is refractory to ICB (*16*), and did not exhibit increased edema under anti-PD1 treatment (**Fig. 2C**).

**Figure 5.**
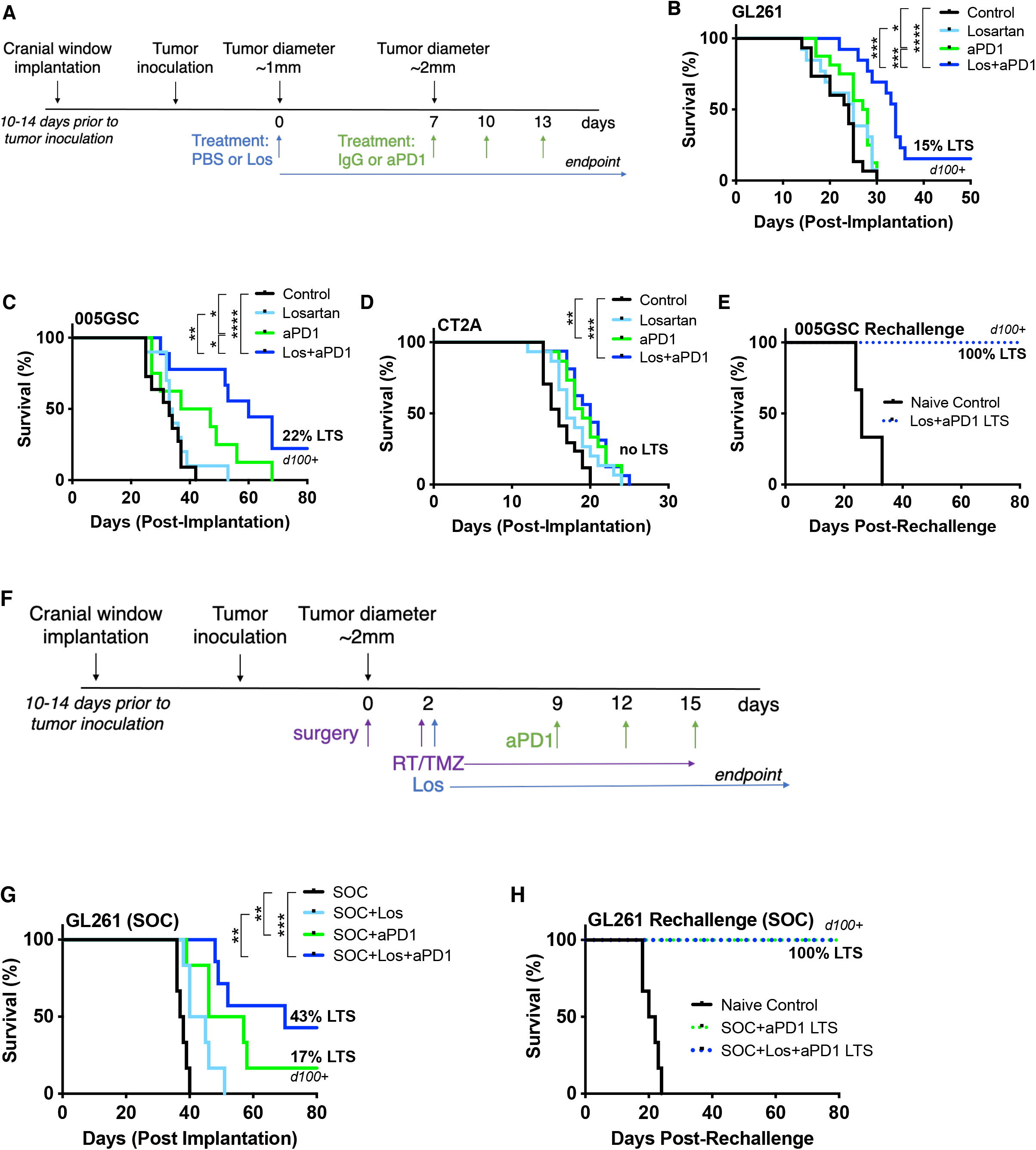
Losartan improves survival under anti-PD1 treatment with and without the standard of care. Losartan enhances the survival benefit of anti-PD1 therapy in (**A**) GL261 and (**B**) 005 GSC tumor models with 15% and 22% long-term survivors (LTS) respectively, with no detectable tumors via microultrasound imaging through transparent cranial windows for over 100 days (d100). In addition to lack of increased edema in the face of ICB treatment (**Fig. 2C**), (**C**) the CT2A model displays only a modest response to anti-PD1 therapy that does not result in long-term survivors nor is improved by the addition of losartan treatment. (**D**) Long-term surviving mice in the 005 GSC model reject a second tumor inoculation, suggesting the formation of an immune memory response. (**E**) The GL261 model subjected to standard of care (SOC; **F**) therapy shows an improvement (**G**) in response to anti-PD1 (16% long-term survivors) that is tripled (43% long-term survivors) in combination with losartan. (**H**) Long-term surviving mice in the GL261 standard of care model reject a second tumor rechallenge. (*Log-rank Mantel-Cox test; * = p<0*.*05; ** = p<0*.*01; *** = p<0*.*001; **** = p<0*.*0001*.)

In GL261 tumors, we found that standard of care treatment (surgical resection, radiation, and temozolomide; **Fig. 5F**) enhanced anti-PD1 outcome to produce 16% long-term survivors (**Fig. 5G**). Long-term survival almost tripled to 43% when losartan was added to standard of care+anti-PD1, and these surviving mice rejected tumor re-challenge (**Fig. 5H**).

### Immune TME biomarkers from bihemispheric tumor model predict individual response to losartan+ICB therapy

Because we observed variable responses in individual mice to losartan+anti-PD1 therapy, we sought to identify predictive biomarkers informed by the GBM immune compartment prior to therapy. Building off our recent bilateral breast cancer model (*17*), we designed a bihemispheric brain tumor model to simultaneously profile immune cells and measure treatment response in individual mice.

Mice were implanted with two identical GL261 tumors in contralateral hemispheres (**Fig. S6**). We resected one tumor for biomarker analysis prior to the initiation of losartan+anti-PD1 therapy. Each resected tumor was profiled for immune cells using flow cytometry. Each mouse (now bearing its remaining non-resected tumor) was evaluated for individual response to losartan+anti-PD1 therapy. Mice were classified based on survival as non-responders, responders (improved median survival), and long-term survivors (no detectable tumor) (**Fig. S6**). We found that, before the initiation of treatment, tumors from long-term surviving mice had strong anti-tumor immune profiles compared to non-responders and responders, including increased ratios of cytotoxic Granzyme B+ CD8 T cells to regulatory FoxP3+ CD4 T cells, and “M1-like” to “M2-like” TAMs and microglia (**Fig. 6A**). Immune biomarkers (T regulatory cells, TAMs, CD4 T cells, and cytotoxic to regulatory T cells ratios) were significantly correlated with survival via univariate Cox proportional hazard models (**Fig. 6B**).

**Figure 6.**
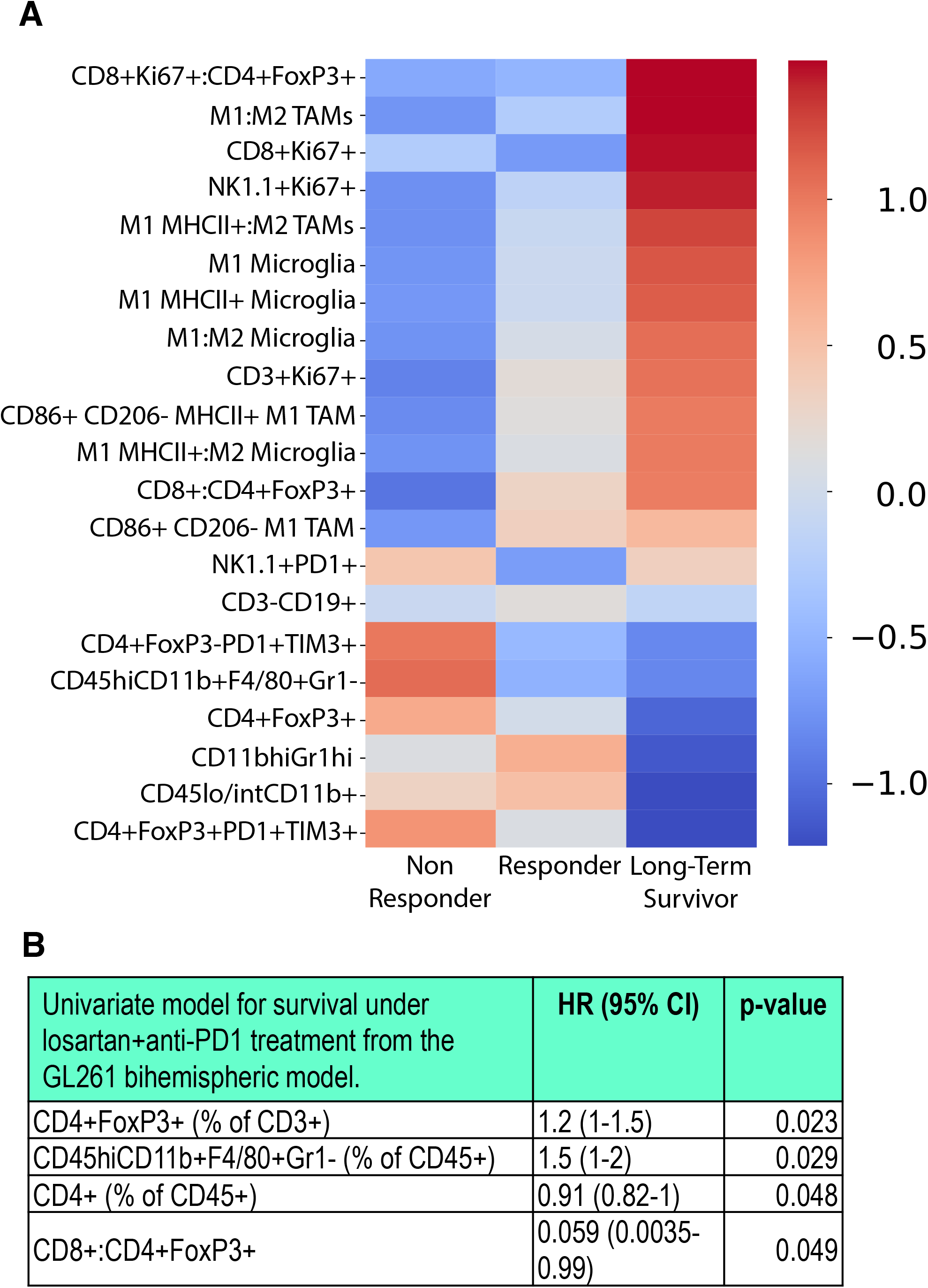
Bihemispheric model reveals predictors of response to losartan+anti-PD1 treatment. The bihemispheric mouse model can be used to resect one tumor for biomarker analysis prior to losartan+anti-PD1 treatment which has variable responses in GL261-bearing mice (n=9). (**A**) Using flow cytometry, immune cells were profiled in individual mice under combinatorial therapy. As indicated by the heat-map *z*-scores (transformed relative populations of immune cell classes), long-term survivors have distinguished pre-treatment biomarker signatures that indicate strong anti-tumor immunity is present in the tumor prior to therapy. (**B**) The presence of CD4 T cells and higher ratios of CD8 to regulatory T cells in the GBM TME before therapy initiation are predictive of improved survival, while the presence of T regulatory cells and TAMs are associated with decreased survival, assessed via proportionate hazard models. (*P-values derived from univariate Cox regression model; HR-Hazard Ratio; CI – Confidence Interval*).

## Discussion

Cerebral edema, a hallmark of GBM, is further exacerbated in a fraction of patients under PD1/PD-L1 treatment (*1, 2*). We sought to identify an agent that could be used in lieu of immunosuppressive corticosteroids – known to compromise ICB efficacy and effector T cell function (*18, 19*) – to control ICB-induced edema.

Losartan is a small molecule ARB commonly prescribed as an anti-hypertensive agent. Losartan can cross the BBB, and ARB use has been reported to be associated with reduced brain edema and lower steroid dosages in GBM patients undergoing chemoradiation treatment (*4, 5, 20, 21*). However, the steroid-sparing edema control mechanism of ARBs is not fully understood. In two syngeneic GBM models, we showed that losartan prevented anti-PD1-induced edema. Brain edema is attributed largely to overexpression of VEGF, which increases vascular permeability (*22*). However, bevacizumab – an anti-VEGF antibody that can control edema – failed to improve OS in GBM patients under ICB therapy (*1*), suggesting a VEGF-independent mechanism for ICB-induced edema.

Our sequencing and T cell blockade studies indicate the involvement of inflammatory edema. Using scRNASeq analysis, we derived a signature of edema prevention in TECs that included downregulation of MT-MMP -1 and -2 by losartan. T cell interactions with endothelial cells increase MMP expression (*8*), which can disrupt tight junctions, leading to a compromised BBB (*6*). However, MT-MMP-1 and 2 have not yet been linked directly to cerebral edema. Our study demonstrates the role of MMPs in mediating anti-PD1-induced edema, generating a working model that CD8+ T cells infiltrating into the GBM TME in response to ICB interact with TECs, inducing their increased expression of MT-MMP-1 and -2. This results in a disrupted blood-tumor-barrier and increased edema. Importantly, although losartan can reduce VEGF (*3*), our results indicate that ICB-induced edema is not VEGF-dependent, but rather due to an inflammatory response.

The immunosuppressive nature of the GBM TME stems from multiple factors. Abnormally high ECM deposition is a key contributor; HA and fibrillar collagens are expressed several-fold higher in GBM than in normal brain tissues (*23, 24*). This contributes to elevated solid stress that impairs perfusion by compressing tumor blood vessels (*25*). Reduced perfusion limits tumor oxygenation, drug delivery, and trafficking of anti-tumor immune cells into the GBM TME. This hostile TME contributes to exclusion and exhaustion of CTLs while promoting the infiltration and activation of immunosuppressive Tregs and pro-tumor myeloid cells including TAMs (*26, 27*). We and others have shown that losartan decreases TGF-β in mice and cancer patients, thus promoting immune stimulation in non-CNS tumors (*11, 12, 28*). However, these effects and the underlying mechanisms have not been investigated in GBM.

Our results indicate that losartan repolarizes TAMs and microglia – both of which promote immunosuppression, and are associated with poor prognosis in GBM (*29*). We recently showed that high expression of the pro-tumor myeloid receptor CCR2 is associated with poor prognosis in GBM patients, and that targeting CCR2 enhances ICB outcome in GBM models (*30*). Our results here indicate that angiotensin inhibition not only reduces the presence of CCR2-positive TAMs and other pro-tumor myeloid cells but also reprograms the compartment to an anti-tumor phenotype. In extracranial mouse and human tumors, we have linked losartan (and similar ARBs) to anti-tumor T cell gene expression, presence and activity (*10, 28, 31*). Here, we observed improved effector T cell infiltration and function during combined losartan+anti-PD1 therapy. Importantly, although losartan reduces inflammatory responses that contribute to ICB-induced edema, it does not abrogate anti-tumor immune activity.

We have shown that losartan (and similar ARBs) can improve response to cytotoxic and ICB in pancreatic and metastatic breast cancer mouse models, respectively (*10, 11*). Here, we found in GBM that losartan improves anti-PD1 outcomes in the 005 GSC and GL261 models, but not in CT2A. This could be due in part to excess ECM deposition in CT2A compared to other models, as well as its lack of responsiveness to ICB, and exclusion and exhaustion of CD8 T cells even in the face of anti-PD1 therapy (*16, 32*). This supposition explains the lack of ICB-induced inflammatory edema in the CT2A model. To lay the groundwork for future clinical translation, we used our recently established standard of care model (*16*) and further improved the durability of losartan+anti-PD1. The lack of secondary tumor formation after re-challenge in “cured” mice suggests the formation of an immune memory response.

Variable patient response to ICB therapy is a stark and challenging clinical reality. There is an unmet need to identify robust and predictive biomarkers of ICB response, due in part to a lack of mechanistic insight into what drives resistance vs. response. This is particularly the case for GBM patients who present with heterogeneous immune landscapes that may drive variable response to ICB (*33-35*). Indeed, we observed differential responses within a single treatment arm, even in genetically-identical mice bearing tumors grown from the same model and batch of GBM cells.

Building on similar approaches in brain, breast, and subcutaneous sites (*17, 36*), we developed a bihemispheric tumor model to predict response to losartan+anti-PD1 immunotherapy. Unlike previous studies, however, we utilized this “resection-and-response” approach to evaluate the composition of the GBM immune compartment *prior* to ICB therapy. Flow cytometry analyses from the bihemispheric model revealed that an immunostimulatory (or “hot”) immune compartment in the TME prior to losartan+anti-PD1 is associated with long-term survivors. This is in line with a recent retrospective transcriptomic analysis showing that patients with “immune-favorable TMEs” benefit the most from immunotherapy (*37*). This approach allows us to establish predictive biomarkers that could be used to inform selection of GBM patients who may respond to losartan+ICB in future clinical trials based on their tumor immune compartment at the time of surgical resection.

A phase III prospective trial with losartan in GBM recently failed to improve median OS in combination with the standard of care (*38*). Similarly, our preclinical results indicate that losartan does not improve OS in GBM mouse models under the standard of care unless it is administered in conjunction with ICB. Recent retrospective studies (e.g., in non-small cell lung, GI, and GU cancers (*39-41*)) suggest that patients under angiotensin system inhibitors may have better response to ICB therapy. Losartan is also under clinical testing for ICB combined with cytotoxic therapy in pancreatic ductal adenocarcinoma patients (NCT03563248) following a successful Phase II trial based on our preclinical findings (*42*). The results of the current study warrant testing combined losartan and ICB therapy in the clinic, along with tissue-based biomarkers identified here for patient selection.

## Acknowledgments

The authors thank the following for their helpful input: Drs. Patrik Andersson, Zohreh Amoozgar, Mark Badeaux, Echoe Bouta, Vikash Chauhan, Jie Chen, Dan Duda, Gino Ferraro, Dai Fukumura, Igor Gomes dos Santos, Peigen Huang, Raymond Huang, Bryan Iorgulescu, Jonas Kloepper, Wilhelmus Kwanten, Louis Larrouquere, Pinji Lei, Hao Liu, John Martin, Hadi Nia, Ethel Periera, Mikael Pittet, Daniel Schanne, Giorgio Seano, Nilesh Talele, Klaus van Leyen, Jan van Wijnbergen, Nancy Wang, Christina Wong. We also thank Anna Khachatryan, Carolyn Smith, and Marni Shore for their technical assistance.

## Funding

National Institutes of Health grant K22-CA258410 (MD); P01-AI56299 (AHS and GJF); P01CA236749 (AHS and GJF); R37-CA245523 (MLS); R35-CA197743 (RKJ); U01-CA224348 (RKJ); R01-CA259253 (RKJ); R01-CA208205 (RKJ); R01-NS118929 (RKJ); U01CA261842 (RKJ).

American Association for Cancer Research-Loxo Oncology postdoctoral fellowship 19-40-50-DATT (MD).

American Brain Tumor Association Basic Research Fellowship (SC).

MGH Fund for Medical Discovery Award (SC).

Pediatric Cancer Research Foundation Young Investigators Award (SC).

National Science Foundation Graduate Research Fellowship Program DGE17453303 (EMP).

Deutsche Forschungsgemeinschaft postdoctoral fellowship DFG, GR 5252/1-1 (SG).

European Union’s Horizon 2020 Programme European Research Council Grant 758657-ImPRESS (KEE).

South-Eastern Norway Regional Health Authority grants 2017073, 2013069 (KEE).

The Research Council of Norway and Cancer Society of Norway grants 261984, 325971, 303249 (KEE).

Agency for Science, Technology and Research (A*STAR) graduate scholarship (WWH, ASK).

Department of Defense Postdoctoral Fellowship W81XWH-19-1-0723 (SK).

Mark Foundation Emerging Leader Award (MLS).

MGH Research Scholars Award (MLS).

Ludwig Cancer Center at Harvard (RKJ).

Nile Albright Research Foundation (RKJ).

Jane’s Trust Foundation (RKJ).

## Author contributions

Conceptualization: MD, AHS, GJF, DAR, MLS, LX, RKJ

Methodology: MD, SC, EMP, SG, SR, MD, IXC, MRN, MCS, MRN, AHS, GJF, DAR, MLS, LX, RKJ

Investigation: MD, SC, EMP, SG, SR, MD, IXC, KN, ASK, MG, KEE, MRN, WWH, PK, SK, XD, MCS, MRN, DAR

Visualization: MD, SC, EMP, SG, IXC, KN, ASK, MG, MCS

Funding acquisition: MD, LX, RKJ

Project administration: MD, LX, RKJ

Supervision: MD, LX, RKJ

Writing – original draft: MD, RKJ

Writing – review & editing: MD, SC, EMP, SG, SR, MD, IXC, KN, ASK, MG, KEE, MRN, WWH, PK, SK, XD, MCS, MRN, AHS, GJF, DAR, MLS, LX, RKJ

## Data and materials availability

Data are available in the supplemental materials (Data S1-S3) or by request.

## Supplementary Materials

Materials and Methods

Figs. S1 to S6

Data S1 to S3 Movies S1 to S2

References (*14-16, 19, 43-48*)

